# Impaired aldehyde detoxification caused by a common human polymorphism promotes anti-bacterial immunity

**DOI:** 10.1101/2023.08.24.554661

**Authors:** Scott Espich, Sam Berry, Amir Balakhmet, Xuling Chang, Nguyen T.T. Thuong, Rajkumar Dorajoo, Chiea-Chuen Khor, Chew-Kiat Heng, Jian-Min Yuan, Woon-Puay Koh, Douglas Fox, Andrea Anaya-Sanchez, Logan Tenney, Christopher J. Chang, Rishika Baral, Dmitri I. Kotov, Marietta Ravesloot-Chavez, Russell E. Vance, Sarah J. Dunstan, K. Heran Darwin, Sarah A. Stanley

**Author notes:** These authors contributed equally to this work (Scott Espich, Sam Berry).

## Abstract

The *ALDH2*2* (rs671) variant present in >500 million individuals reduces ALDH2 function, impairing aldehyde detoxification. While aldehyde accumulation in these individuals is associated with numerous negative health consequences, a previous study showed a cohort of *ALDH2*2* carriers are less likely to develop active pulmonary tuberculosis. Here, we present additional human data that support this finding and show ALDH2-deficiency in mice provides a fitness advantage during bacterial infections. We found aldehydes normally detoxified by ALDH2 killed the bacterial pathogens *Mycobacterium tuberculosis* and *Francisella tularensis.* Infected macrophages from *Aldh2^−/−^*mice had higher levels of formaldehyde and 4-hydroxynonenal, which enhanced their microbicidal capacity. *Aldh2^−/−^* mice were more resistant to infection with *Mycobacterium tuberculosis* and *Francisella tularensis* than parental mice and displayed elevated inflammatory cytokine and chemokine levels, accompanied by an increased accumulation of inflammatory monocytes and macrophages. These findings support a model in which host-derived aldehydes are robust innate immune effectors, limiting bacterial infection through both direct microbicidal activity and immune modulation. Collectively, this work may explain why the *ALDH2*2* allele was selected for in humans.

## INTRODUCTION

Loss-of-function mutations in aldehyde dehydrogenase 2 (*ALDH2*) are highly prevalent, affecting up to 500 million individuals of East Asian descent^1^. The most common mutant allele, *ALDH2*2* (rs671), encodes a glutamate-to-lysine mutation at position 487, resulting in a dominant-negative form of the enzyme^2^. ALDH2 is notably required for converting acetaldehyde, a toxic intermediate of ethanol metabolism, into acetate, but its primary function is to detoxify metabolically produced aldehydes. Hetero- or homozygous carriers of *ALDH2*2* experience adverse physiological effects upon consumption of alcohol due to an accumulation of acetaldehyde^3^. Even in the absence of alcohol consumption, loss of ALDH2 function is associated with increased risk of cancer, heart disease, and neurological disorders^4,5^. Aldehydes can react with DNA, leading to mutations, and may disrupt cellular function by forming adducts with proteins, contributing to disease development^6–8^. Additionally, loss of ALDH2 activity has been associated with heightened inflammatory responses, which may further promote disease pathogenesis^9,10^. The selective pressure that has maintained *ALDH2*2* in the human population despite its detrimental effects on human health is unknown.

*ALDH2*2* was associated with a decreased risk of active tuberculosis (TB) in a cohort of Korean men. This correlation was hypothesized to be associated with decreased alcohol consumption^11^. While alcoholism is a significant risk factor for TB^12,13^, we hypothesized that loss of ALDH2 function enhances resistance to bacterial infection independent of alcohol consumption. ALDH2 detoxifies many inflammation-induced aldehydes, including 4-hydroxynonenal (4HNE)^14–17^, a lipid peroxidation end-product that is produced when macrophages and other innate immune cells generate reactive oxygen species in response to infection^16,18–20^. Additionally, ALDH2, along with alcohol dehydrogenase 5 (ADH5), metabolizes endogenous formaldehyde, which is produced by a range of cellular processes^21,22^. Formaldehyde production is also enhanced in the presence reactive oxygen species^23^. Together these studies suggest that loss of ALDH2 activity during infections could increase cellular aldehyde levels that would be toxic to invading bacteria. Here, we demonstrate that loss of ALDH2 function enhances control of bacterial infection by promoting the microbicidal activity of macrophages and by accelerating inflammatory responses to infection.

## RESULTS

### ALDH2 rs671 is associated with lower pulmonary tuberculosis risk

Pulmonary tuberculosis (PTB) is the most common clinical manifestation of TB disease and leads to the principal mode of transmission amongst humans. To examine the relationship between the *ALDH2*2* mutation (rs671) and PTB, we performed a candidate gene analysis for cohorts in Vietnam and Singapore. Genotype data of rs671 was available from a collection of 1,598 microbiologically confirmed TB patients and 1,266 population controls from Vietnam (Figure 1A). In addition, the Singapore Chinese Health Study (SCHS) consists of 1,606 PTB cases and 23,969 population controls of Singaporean Chinese origin with available genotype data. A significant association between rs671 PTB was observed in both the Vietnamese cohort [OR (95% CI) = 0.855 (0.739, 0.990), *P*=0.036] and the SCHS [OR (95% CI) = 0.920 (0.850, 0.997), *P*=0.041] (Figure 1B), with the minor allele A associated with reduced PTB risk. After meta-analysis, the association was further strengthened ([OR (95% CI) = 0.905 (0.844, 0.971), *P*=0.005, *P*_heterogeneity_=0.386]). We also assessed both dominant and recessive models of inheritance and found the dominant model returned associations of similar magnitude to the additive model [OR (95% CI) = 0.888 (0.813, 0.969), *P*=0.008, *P*_heterogeneity_=0.429]. The modest effect sizes calculated, which were robustly replicated across two ethnic groups, are not unexpected, as the heritability of complex traits, such as infectious diseases, arises from the combined influence of numerous genetic variants, each contributing a modest effect size. These data, together with the previously published association study^11^, support the hypothesis that the *ALDH2*2* rs671 mutation is associated with protection against PTB.

**Figure 1.**
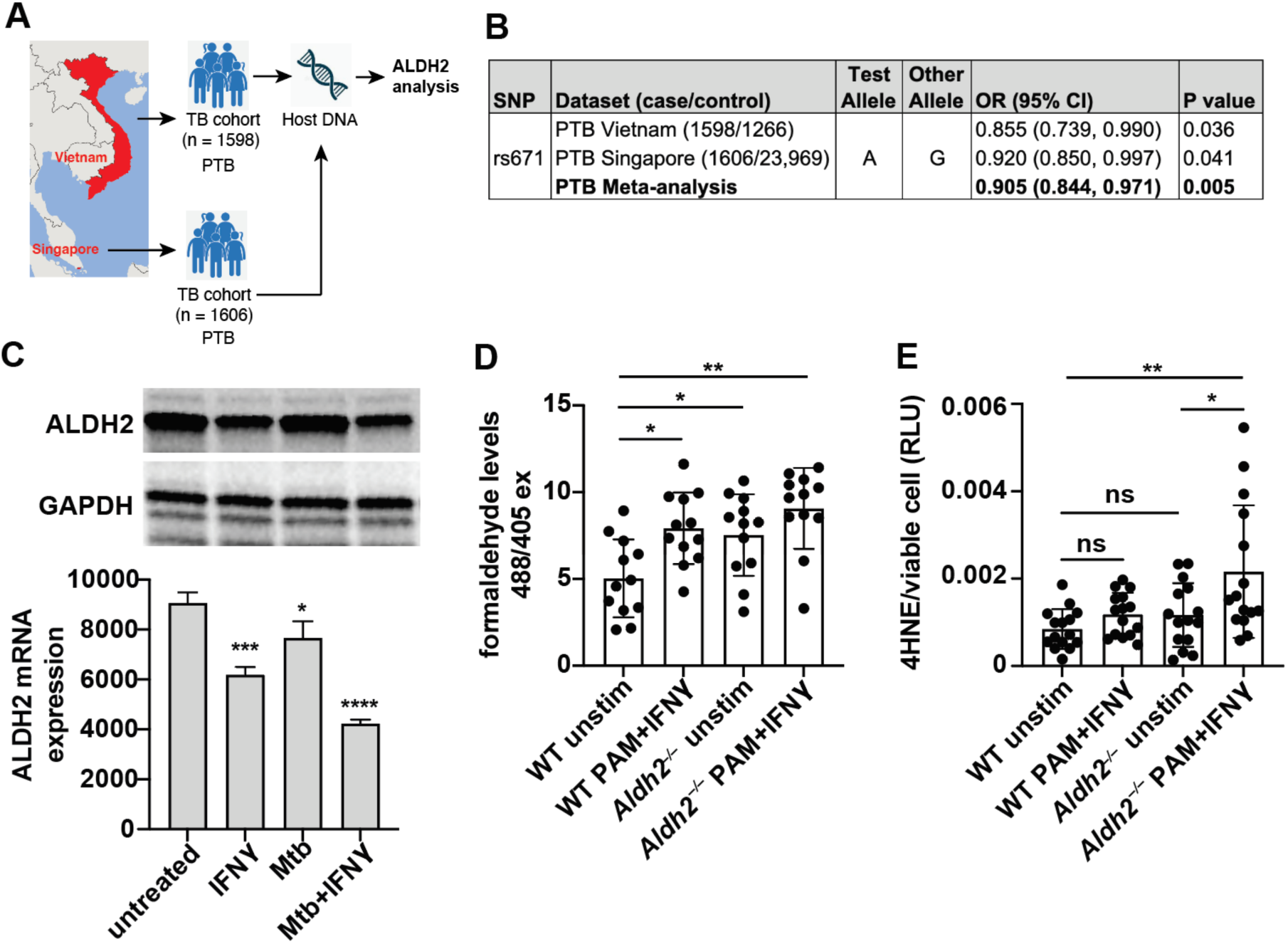
ALDH2 expression is linked with human disease and immune activation. (A, B) A candidate gene analysis of the ALDH2 rs671 mutation in pulmonary TB (PTB) patients in Vietnam and Singapore. (C) mRNA and protein for *Aldh2* expression *M. tuberculosis* infected C57BL/6J (WT) bone marrow-derived macrophages (BMDMs) stimulated with IFNγ and/or infected with *M. tuberculosis* H37Rv. (D) WT and *Aldh2^−/−^* BMDM mitochondrial-associated formaldehyde levels after 24 hours stimulation with IFNγ and PAM3CSK4 *in vitro.* (E) ELISA quantification of BMDM 4HNE after 24 hours stimulation with IFNγ and PAM3CSK4 *in vitro.* Experiments are representative of three independent experiments (C, D) or three independent experiments combined (E). (C, D) unpaired t-test, (E) Two-way ANOVA with Tukey’s multiple comparisons test. *, *p* < 0.05, **, *p* < 0.01, ***, *p* < 0.001, ****, *p* < 0.0001.

### *Aldh2* expression in macrophages is downregulated upon inflammatory stimulation

To determine whether ALDH2 is expressed in relevant cell types during infection, we examined *Aldh2* expression in the lungs of *M. tuberculosis* infected C57BL/6J (“WT”) mice using single cell RNA sequencing (scRNAseq). *Aldh2* expression was observed in macrophage/monocyte populations, dendritic cells, neutrophils, basophils, B cells, pulmonary epithelial and endothelial cells, and fibroblasts (Figure S1A). Other immune cell types, such as T cells, did not express *Aldh2*. Macrophages serve both as the primary host cell for *M. tuberculosis* and a key effector cell in infection control, so we next investigated how infection alters *Aldh2* expression in bone marrow–derived macrophages (BMDMs) from WT mice. We infected BMDMs in the presence or absence of interferon-γ (IFNγ), a critical cytokine for macrophage activation and control of *M. tuberculosis* infection^24^. Consistent with the *in vivo* scRNA-seq data, ALDH2 protein was expressed in unstimulated BMDMs (Figure 1C). In contrast, IFNγ stimulation reduced *Aldh2* expression in both uninfected and *M. tuberculosis*-infected macrophages (Figure 1C), suggesting that IFNγ induces a transcriptional program that downregulates *Aldh2* expression during infection.

### Macrophages from *Aldh2*-deficient mice produce higher aldehyde levels

We next generated *Aldh2* mutant (*Aldh2*^−/−^) mice using CRISPR-Cas9 (Figure S1B). The mutation (12 base pair deletion in exon 12 that encompasses the region where rs671 is found in humans) resulted in loss of ALDH2 as determined by immunoblotting (Figure S1C). We tested if immune activation could increase aldehyde production by wild-type (WT) macrophages and whether aldehyde production would be further enhanced by *Aldh2*-deficiency (Figure S1D). Using Mito-RFAP-2, a mitochondrial-targeted, ratio-metric dye that detects mitochondrial-localized formaldehyde with high sensitivity^25^, we found that WT macrophages stimulated with IFNγ and PAM3CSK4, a TLR2 agonist, had increased formaldehyde 24 hours after stimulation (Figure 1D). This stimulation-induced enhancement of aldehyde production in macrophages inversely correlated with *Aldh2* expression (Figure 1C) and increased mitochondrial formaldehyde levels (Figure 1D). *Aldh2^−/−^* macrophages produced significantly more formaldehyde than the WT cells in the absence of immune stimulation (Figure 1D). A significant difference was not observed between stimulated and unstimulated *Aldh2^−/−^* macrophages, suggesting that formaldehyde levels in *Aldh2^−/−^* macrophages could not be further increased by external stimulation.

We next measured 4HNE-protein adducts by ELISA. Although we observed a trend towards increased 4HNE production in response to stimulation with PAM3CSK4 and IFNγ in WT macrophages, it was not significantly different between stimulated and resting macrophages (Figure 1E). However, stimulated *Aldh2^−/−^* macrophages produced higher levels of 4HNE than stimulated WT or unstimulated *Aldh2^−/−^* macrophages (Figure 1E).

### Aldehydes detoxified by ALDH2 kill *M. tuberculosis*

Based on the role of ALDH2 in detoxifying inflammation-associated aldehydes, we tested the susceptibility of *M. tuberculosis* to 4HNE and formaldehyde in axenic culture. 4HNE can reach millimolar concentrations in inflamed tissues^8,15,26^, and formaldehyde has been found at up to 400 *μ*M in blood and tissues^27–29^. Using a standard half-maximal inhibitory concentration (IC_50_) protocol, we found the *in vitro* IC_50_ values range between 40-150 µM for formaldehyde while 4HNE had an IC_50_ of 40 µM. *M. tuberculosis* was sensitive to 4HNE, with 10-fold killing observed at 50 *μ*M after six days of exposure and greater than three orders of magnitude of toxicity at 200 *μ*M (Figure 2A) relative to controls. Similarly, we found that formaldehyde was toxic to *M. tuberculosis* at 100 *μ*M (Figure 2B). Thus, *M. tuberculosis* is susceptible to physiologically relevant levels of aldehyde in broth culture.

**Figure 2.**
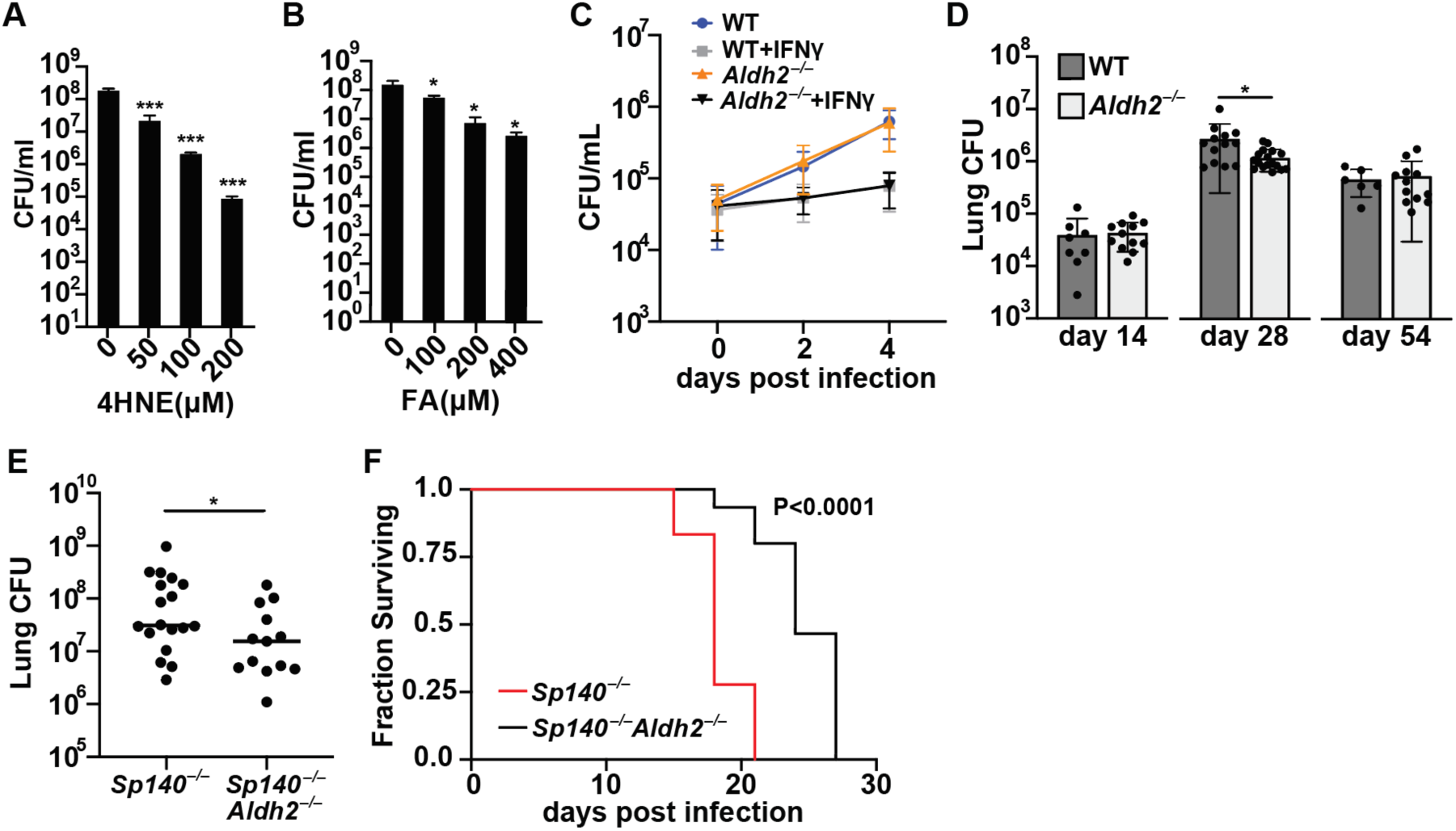
*Aldh2*-deficient mice exhibit enhanced control of *M. tuberculosis* infection. *M. tuberculosis* H37Rv killing after six days treatment with physiological levels of (A) 4-hydroxynonenal (4HNE) or (B) Formaldehyde (FA). (C) CFU enumeration of WT and *Aldh2^−/−^*BMDMs infected with *M. tuberculosis* LuxErd +/− IFNγ at an MOI= 1. (D) Lung CFU enumeration of aerosol-infected *Aldh2^−/−^* and WT mice with ≈ 25 CFU *M. tuberculosis* H37Rv at indicated timepoints. (E) Lung CFU enumeration of aerosol-infected *Sp140^−/−^* and *Sp140^−/−^Aldh2^−/−^* mice with ≈ 25 CFU *M. tuberculosis* Erdman 28 dpi. (F) Survival curve of *Sp140^−/−^* and *Sp140^−/−^Aldh2^−/−^* mice infected with a high dose of *M. tuberculosis* Erdman (300 CFU). (A), (B) *p<0.05 by unpaired t-test. (D), (E) p<0.05 by Mann-Whitney. (F) p<0.0001 by Mantel-Cox log rank test. Experiments are representative of a minimum of three (A), (B), (C), (F) or three experiments combined (D), (E).

### Mice lacking *Aldh2* exhibit reduced susceptibility to *M. tuberculosis* infection

We next tested whether ALDH2 deficiency enhances control of *M. tuberculosis* by infected macrophages. WT and *Aldh2*^−/−^ macrophages were infected with *M. tuberculosis* at a multiplicity of infection of one bacillus per macrophage (MOI = 1), and bacterial replication was monitored over four days. *M. tuberculosis* replication was identical in WT and *Aldh2*^−/−^ macrophages (Figure 2C). We observed the expected restriction of *M. tuberculosis* growth after IFNγ activation, but we did not observe significant differences between the WT and *Aldh2*^−/−^ BMDMs (Figure 2C). Given that *in vitro* conditions neither recapitulate the extent of inflammation nor the complete pathogenesis observed *in vivo*, we infected *Aldh2^−/−^* mice to assess the role of ALDH2 in a biologically relevant model of *M. tuberculosis.* We infected mice via the aerosol route with a low-to-medium dose (∼25 colony forming units, CFU) of *M. tuberculosis* and harvested their lungs for CFU enumeration at various time points. At 14 days post infection (dpi), we observed no difference in the ability of *Aldh2^−/−^* mice to control pulmonary infection when compared with littermate controls (Figure 2D); however, at 28 dpi, we observed a significant difference between *Aldh2^−/−^* mice and WT mice, wherein *Aldh2*^−/−^ mice had lower bacterial burdens (Figure 2D). The adaptive immune response is typically observed after 14 dpi in mice, where increased IFNγ signaling results from T cell migration to the site of infection. This result suggests that sustained T cell-mediated IFNγ signaling may contribute to the improved control observed by the *Aldh2^−/−^* mice at 28 dpi by increasing local aldehyde production. WT mice are relatively resistant to infection with *M. tuberculosis*^30,31^.

We hypothesized that the protective effect of ALDH2 could be more robust in a susceptible mouse background in which a loss of bacterial control correlates with destructive inflammatory responses in the lung, as well as in a setting of increased aldehyde load. To test this hypothesis, we generated *Aldh^−/−^* mice on the background of *Sp140* deficiency. *Sp140^−/−^* mice have type I IFN-driven susceptibility to disease and severe bacterial burden during *M. tuberculosis* infection^32^. We infected *Aldh2^−/−^ Sp140^−/−^* and parental *Sp140^−/−^*mice with *M. tuberculosis*. Both genotypes had increased lung CFU relative to *Aldh2^−/−^* and WT mice due to the known susceptibility of the *Sp140^−/−^* background (Figure 2E). As with WT mice, we observed significantly fewer CFU in the *Aldh2^−/−^Sp140^−/−^* mice when compared to parental *Sp140^−/−^* animals at 28 dpi (Figure 2E). To determine if *Aldh2* deficiency promotes survival during infection in this context, we infected *Aldh2^−/−^Sp140^−/−^*mice with a high dose of *M. tuberculosis. Aldh2^−/−^Sp140^−/−^* mice lived significantly longer that the *Sp140^−/−^*controls (Figure 2F). Thus, even in the presence of a host-detrimental inflammatory type I IFN response, disruption of *Aldh2* had a protective effect against *M. tuberculosis* infection.

### *Aldh2* deficiency enhances control of *Francisella* infection

Previous work demonstrated the susceptibility of *Francisella* species to 4HNE and other metabolic aldehydes^33,34^. We observed that the *Francisella tularensis* Live Vaccine Strain (FtLVS) was highly susceptible to 4HNE and formaldehyde, showing nearly three orders of magnitude of killing with 50 μM 4HNE, and ∼1 log killing with 25 μM formaldehyde (Figures 3A and 3B). To test whether *Aldh2^−/−^* mice are resistant to infection with FtLVS, we infected mice with FtLVS intradermally (i.d.) and measured CFU in lungs and spleens five days later. We found that in both organs, *Aldh2^−/−^* mice had significantly lower bacterial burdens than parental mice (Figures 3C and 3D). We also observed elevated levels of 4HNE concentrations in tissues of infected *Aldh2^−/−^* mice (Figures 3E and 3F). Because macrophages are a primary replicative niche for FtLVS^35^, we next reasoned that enhanced control of infection in *Aldh2^−/−^* mice could be associated with enhanced macrophage microbicidal activity. To test this hypothesis, we infected BMDMs from WT and *Aldh2^−/−^* mice *in vitro* and found that *Aldh2^−/−^* macrophages were indeed more microbicidal than WT macrophages, an effect that was enhanced by IFNγ stimulation (Figure 3G). CFU was equivalent across genotypes on day 0, and no differences in macrophage viability were observed at any timepoint (Figures S3J and S3K).

**Figure 3.**
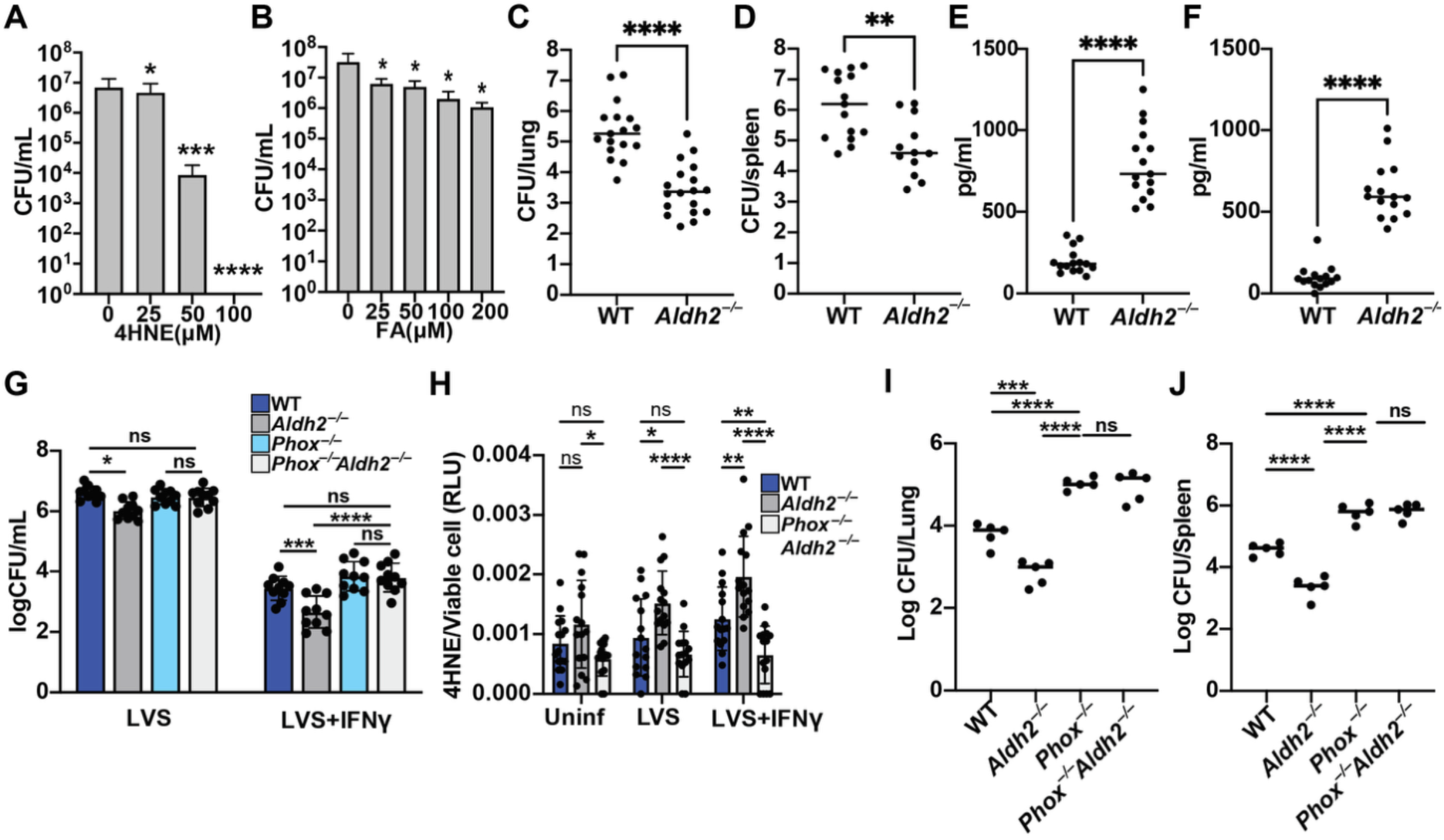
*F. tularensis* LVS growth is attenuated in *Aldh2^−/−^* mice and macrophages. *F. tularensis* LVS (FtLVS) killing after two hours of treatment with physiological levels of (A) 4HNE or (B) formaldehyde. CFU enumeration of WT and *Aldh2^−/−^* lungs (C) or spleens (D) infected with 10^5^ FtLVS i.d., five dpi. 4HNE quantification in WT and *Aldh2^−/−^* lungs (E) or spleens (F) infected with 10^5^ FtLVS i.d., 3 dpi. (G) CFU enumeration of WT and *Aldh2^−/−^* BMDMs infected with FtLVS +/− IFNγ at an MOI= 1. (H) 4HNE quantification in untreated or FtLVS infected BMDMs +/− IFNγ. FtLVS CFU enumeration in (I) lungs or (J) spleens of WT, *Aldh2^−/−^, p47phox^−/−^* and *Aldh2^−/−^ p47phox^−/−^* mice infected with 10^3^ FtLVS i.d., 5 dpi. *p<0.05, **p<0.01, ***p<0.001, ****p<0.0001. (A-D) unpaired t-test. (E, F) Mann-Whitney. (G, H) 2way ANOVA with Tukey’s multiple comparisons test. (I-J) one-way ANOVA with Dunnett’s multiple comparisons test. Experiments are representative of a minimum of three (A), (B), (I), (J) or three experiments combined (C-H).

To further correlate the microbicidal activity of *Aldh2^−/−^* macrophages with elevated aldehyde levels, we sought to abolish infection-associated aldehyde production in *Aldh2^−/−^* macrophages and mice. Lipid peroxidation reactions downstream of NADPH oxidase are the primary source of 4HNE in activated macrophages^36^. Consistent with this observation, we found that macrophages lacking both *Aldh2* and *p47phox* (*phox^−/−^*), a subunit of NADPH oxidase, had WT 4HNE levels after infection with FtLVS (Figure 3H). Consistent with the lowered aldehyde levels, loss of *p47phox* also abolished the enhanced control of infection observed in *Aldh2^−/−^* macrophages (Figure 3G). Deletion of *p47phox* also reversed *Aldh2* dependent control of FtLVS infection in vivo (Figures 3I and 3J). We were unable to consistently detect formaldehyde in FtLVS infected macrophages.

### *Aldh2^−/−^* mice have enhanced innate responses to FtLVS infection

To test whether differences in inflammatory responses contribute to enhanced FtLVS control in *Aldh2^−/−^* mice, we analyzed pulmonary and splenic immune responses three dpi., a timepoint preceding significant differences in bacterial burden (Figures 4A and 4B). We found that *Aldh2^−/−^* mice showed a selective increase in recruitment of inflammatory monocytes (iMOS) and inducible nitric oxide synthase (iNOS)-expressing macrophages relative to WT controls. Frequencies of neutrophils and other cell types remained comparable between genotypes (Figures 4C-H and S4). Further, we found that Chemokine (C-C motif) ligand 2 (CCL2) levels were elevated in both the lungs and spleens of *Aldh2^−/−^* mice, explaining the increased recruitment of monocytes and macrophages (Figures 4I and 4J). In addition to CCL2, we found that numerous inflammatory cytokines were elevated in *Aldh2^−/−^* spleens and lungs (Figures S3A-I). Although iNOS^+^ macrophages were enriched in the spleen and lungs of *Aldh2^−/−^* mice, protection was maintained in the absence of iNOS, indicating that aldehyde-mediated control is independent of NO production under these conditions (Figures 4K and 4L).

**Figure 4.**
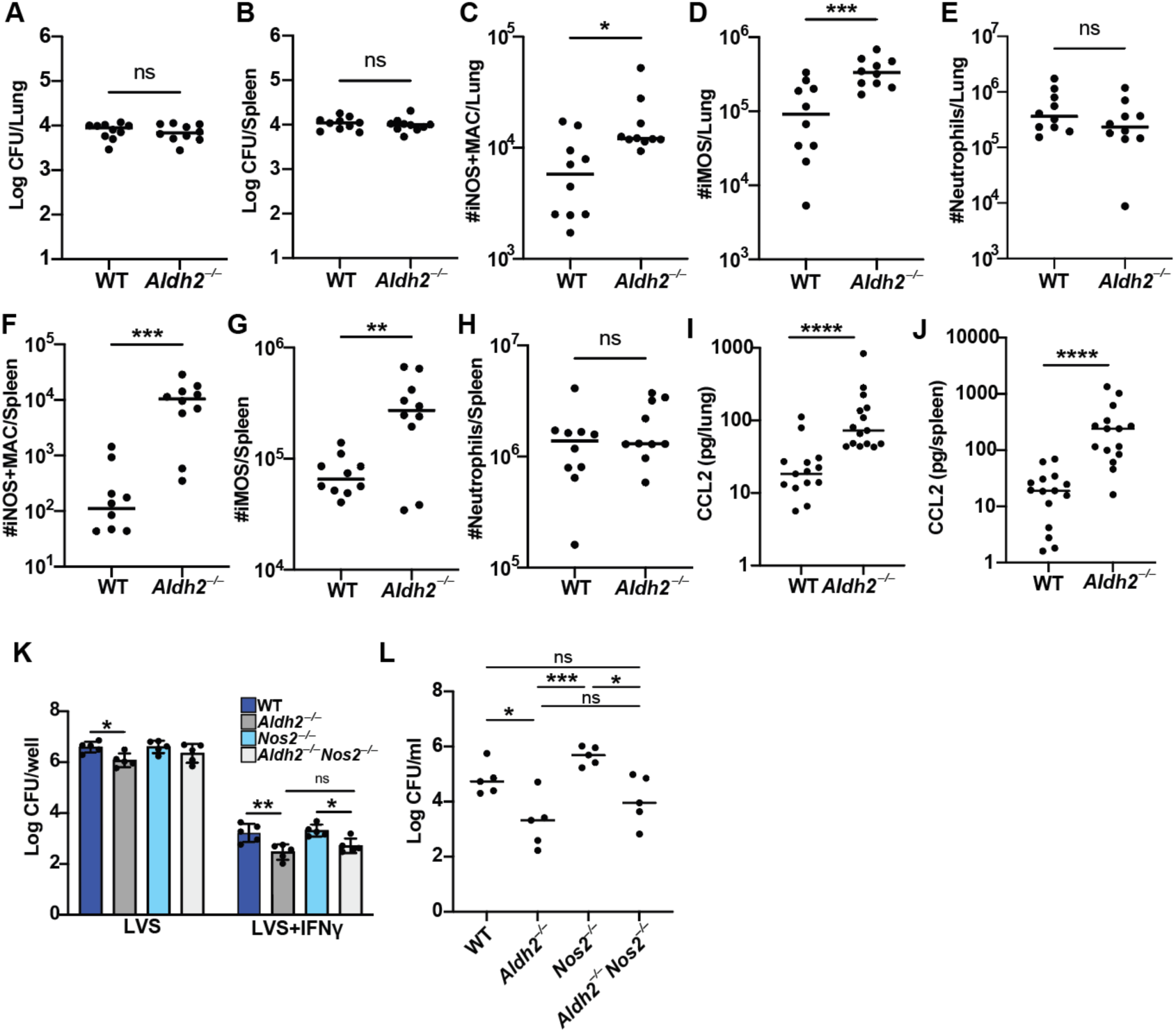
*Aldh2^−/−^*mice exhibit heightened innate immune activation following *F. tularensis* LVS infection. CFU enumeration of lung (A) and spleen (B) in WT and *Aldh2^−/−^* mice infected with 10^5^ FtLVS i.d., 3 dpi. Frequencies of iNOS^+^ macrophages (C) inflammatory monocytes (D) and neutrophils (E) in the lungs of WT and *Aldh2^−/−^* mice infected with 10^5^ FtLVS i.d, 3 dpi. Frequencies of iNOS^+^ macrophages (F) inflammatory monocytes (G) and neutrophils (H) in the spleens of WT and *Aldh2^−/−^* mice infected with 10^5^ FtLVS i.d., 3 dpi. CCL2 quantification in the lungs (I) and spleens (J) of WT and *Aldh2^−/−^* mice infected with 10^5^ FtLVS i.d., 3 dpi. (K) CFU enumeration in WT, *Aldh2^−/−^, Nos2^−/−^,* and *Nos2^−/−^Aldh2^−/−^* infected with FtLVS MOI=1, +/− IFNγ, 2 dpi. L) CFU enumeration in WT, *Aldh2^−/−^,Nos2^−/−^,* and *Nos2^−/−^Aldh2^−/−^*mice infected with 10^3^ FtLVS, 5 dpi. *p<0.05, **p<0.01, ***p<0.001, ****p<0.0001. (A), (B): unpaired t-test. (C-J): Mann-Whitney test. (K): 2way ANOVA with Tukey’s multiple comparisons test. (L): one-way ANOVA with Dunnett’s multiple comparisons test. Experiments are representative of a minimum of three (A-J), (K), (L) or three experiments combined (I), (J).

## DISCUSSION

Throughout human history, pathogens have acted as powerful selective forces, shaping the genetic architecture of modern populations. Classic examples, such as the persistence of the sickle cell allele in malaria endemic regions, illustrate how genetic variants that confer resistance to pathogens can be maintained despite associated fitness costs. In this context, our findings suggest that the widespread ALDH2*2 allele, though traditionally associated with alcohol intolerance, may represent an evolutionary adaptation to bacterial infection^37^. We show that ALDH2 deficiency confers enhanced protection against *Mycobacterium tuberculosis* and *Francisella tularensis* LVS, both in humans (for Mtb) and in mouse models (both pathogens). Mechanistically, *Aldh2^−/−^* mice and macrophages have elevated levels of bactericidal 4HNE and formaldehyde, enhanced innate immune activation, and increased bacterial killing. Taken together, our results support a model in which loss of ALDH2 function amplifies aldehyde-mediated antimicrobial activity, potentially explaining the positive selection of *ALDH2*2* in pathogen-exposed populations. This may represent a previously unrecognized instance of balancing selection, in which antagonistic pleiotropy maintains *ALDH2*2*, despite its deleterious effects, by enhancing protection against infection.

The *ALDH2*2* allele is predicted to have evolved about 3,000 years ago^38^ and is predominantly found in individuals of east Asian descent; interestingly, the emergence of *M. tuberculosis* lineage 2 (also known as the W/Beijing lineage) occurred between 2,000 – 7,000 years ago^38,39^, and this lineage is more prevalent in East Asia. In congruence with previous data^11^, we found that individuals harboring the *ALDH2*2* allele have a reduced risk of developing PTB, suggesting that there may have been evolutionary pressure selecting for this allele in environments where TB transmission was high. Lineage 2 strains exhibit increased transmissibility, hypervirulence, and a higher mutation rate^40–43^, all of which may facilitate survival under host-imposed antimicrobial pressures—including aldehyde stress. Further research is needed to explore if there is an evolutionary interplay between *ALDH2*2* and lineage 2 strains.

Although *Aldh2^−/−^* mice showed early protection against *M. tuberculosis,* we did not observe differences in bacterial burden at later timepoints. One explanation is that in the relatively resistant B6 background, infection becomes persistent and controlled by 28-56 dpi, with limited bacterial replication and dampened inflammation, conditions under which aldehyde-mediated killing may be less effective. In contrast, *Aldh2* deficiency confers an advantage in *Sp140^−/−^*mice, even when infection is acute and poorly controlled. Consistent with this result, we observed robust protection in *Aldh2^−/−^* mice challenged with FtLVS, a model of acute, rapidly progressive infection. Importantly, our finding that individuals carrying the *ALDH2*2* allele have reduced risk of PTB suggests that even modest enhancements in early host control may confer meaningful protection in natural infection settings.

ALDH2 deficiency has been associated with heightened inflammation in both clinical and experimental settings^10,44^. The accumulation of reactive aldehydes can act as an endogenous danger signal by modifying host proteins and activating stress and immune signaling pathways^45^. Our findings extend this connection to the context of infection. Notably, protection in *Aldh2^−/−^*mice and macrophages was dependent on NADPH oxidase, which drives reactive oxygen species production within the phagosome following bacterial phagocytosis. This result suggests that aldehydes produced via phagosomal lipid peroxidation reactions are required for protection observed in *Aldh2^−/−^*mice. Further, flow cytometry analysis revealed that *Aldh2^−/−^*mice infected with FtLVS exhibited increased recruitment of inflammatory monocytes and macrophages to infected tissues, which was accompanied by elevated levels CCL2 and IFNγ, a potent inflammatory macrophage activator required to protect against FtLVS infection. These data suggest that NADPH oxidase-driven aldehyde accumulation not only contributes to direct bacterial killing but may also amplify innate immune activation that promotes pathogen clearance. This idea is consistent with previous studies that show that *ALDH2* deficiency is associated with increased frequencies of inflammatory macrophages^46^.

FtLVS evades early innate immune recognition, a strategy critical for its pathogenesis^47^. Aldehyde-induced early inflammation may disrupt this evasion, restricting bacterial replication and partially attenuating infection. Supporting this hypothesis, pharmacological inhibition of ALDH2 increases macrophage resistance to FtLVS infection *in vitro,* suggesting that aldehyde accumulation alone may be sufficient to enhance antibacterial immunity^48^. While our study focused on 4HNE and formaldehyde, ALDH2 detoxifies a broad spectrum of endogenous aldehydes, including malondialdehyde and acrolein, which may also accumulate in *Aldh2*^−/−^ mice and contribute to host protection. Future work will define the full repertoire of immunomodulatory and antimicrobial aldehydes regulated by ALDH2, and how their cumulative effects shape the immune response to infection.

Collectively, our findings highlight the dual role of aldehydes as both cytotoxic metabolites and immune effectors^37^. More broadly, this work underscores the importance of metabolic regulation in innate immunity and suggests that host-derived aldehydes represent a novel class of endogenous antimicrobial factors weaponized by host cells to combat pathogens. Elucidating how these reactive metabolites function in different infection models and inflammatory contexts may open new avenues for host-directed therapeutic strategies.

## RESOURCE AVAILABILITY

### Lead contact

Further information and requests for resources and reagents should be directed to, and will be fulfilled by, the lead contact, Sarah A. Stanley.

### Materials availability

All reagents generated or used in this study are available on request from the lead contact with a completed materials transfer agreement.

### Data and code availability

All data reported in this paper will be shared by the lead contact upon request. Any additional information required to reanalyze the data reported in this work paper is available from the lead contact upon request.

## Acknowledgements

We thank Angus Y. Lee and Harmandeep S. Dhaliwal (UC Berkeley Cancer Research Laboratory) for generation of *Aldh2* mutant mice. We thank all the participants involved in the genetics studies in Vietnam. and Singapore. We acknowledge the clinical and laboratory staff at the study sites in Vietnam and Singapore for their contribution, especially Phan Vuong Khac Thai, Maxine Caws, Nguyen Duc Bang, Tran Thi Hong Chau, Guy Thwaites, and their teams at Oxford University Clinical Research Unit, and Pham Ngoc Thach Hospital for TB and Lung Diseases, Ho Chi Minh City, Vietnam. We thank Zheng Li from the Genome Institute of Singapore for his contribute to the TB GWAS. We thank Karen Elkins (FDA) for the gift of the Francisella LVS strain as well as helpful advice and protocols.

## Author contributions

Conceptualization: KHD, SAS; Data Curation: SAS, KHD, SE, SB; Formal analysis: SE, SB, AB, XC, DK, DF, SAS; Investigation: SE, SB, AB, RB, DF, LT, AAS, LT; Resources: NTTT, RD, CCK, CKH, JMY, WPK, CJC, LT; Supervision: RV, SJD, SAS, KHD, CJC; Writing: SAS, SE, SJD; Review and Editing: all authors.

## Materials and Methods

### Ethics Statement

All procedures involving the use of mice were approved by the University of California (UC) Berkeley, Institutional Animal Care and Use Committee (Protocol #2015-09-7979-1). All protocols conform to federal regulations, the National Research Council Guide for the Care and Use of Laboratory Animals, and the Public Health Service Policy on Humane Care and Use of Laboratory Animals. All experiments handling *M. tuberculosis* were conducted in Biosafety Level 3 facilities in accordance with Environmental Health and Safety guidelines, and all experiments handling *F. tularensis* LVS were conducted in Biosafety Level 2 facilities. Study protocols for PTB human studies were approved by the Institutional Review Boards of Pham Ngoc Thach Hospital for TB and Lung Disease and The Hospital for Tropical Diseases Ho Chi Minh City (HCMC), Vietnam, HCMC Health Services, Vietnam and the Oxford Tropical Research Ethics Committee, UK. Approval for the genetics study was also granted from the Health Sciences Human Ethics Sub-Committee at the University of Melbourne, Australia (ID: 1340458). Written informed consent was obtained from all participants. All studies in SCHS were approved by the Institutional Review Board at National University of Singapore, and written informed consent was obtained from all participants.

### Mice

C57BL/6J (#000664), *Nos2^−/−^,* and p*47phox^−/−^* mice were obtained from The Jackson Laboratory (Bar Harbor, ME) and bred in-house. *Sp140^−/−^* mice were generated previously^32^ and bred in-house. *Aldh2^−/−^* mice were generated in the process of recapitulating the human *ALDH2*2* mutation in *Sp140^−/−^* mice. *Aldh2*2Sp140^−/−^* mice were generated by pronuclear injection of Cas9 mRNA, sgRNA, and a 154 nucleotide ultramer oligo (IDT Inc., Coralville, IA) containing 77 and 39 nucleotide homology arms and encoding the *Aldh2*2* mutation as well as synonymous mutations that disrupt sgRNA binding, as previously described^49^. Founder mice were genotyped for *Aldh2*2* mutations by PCR using the forward primer 5’ GGAGAGGTGTGCTGTGTT 3’ and the reverse primer 5’ GGGATCTGTATTCCGTGGC 3’ and then sequencing using the same forward primer. Founders carrying the *Aldh2*2* mutation or deletions in *Aldh2* (i.e., *Aldh2^−/−^*) were bred one generation to *Sp140^−/−^, Nos2^−/−^,* or p47*phox^−/−^* mice to separate modified *Aldh2* haplotypes. Resulting pups with matched *Aldh2* haplotypes were interbred to generate homozygous lines that were maintained in-house.

### Bacterial Culture

Frozen stocks of low passage *M. tuberculosis* H37Rv, *M. tuberculosis* Erdman, and *M. tuberculosis* Lux-Erdman (a reporter strain that constitutively expresses the bacterial luciferase encoding luxCDABE operon, gift from Cox Laboratory at UC Berkeley) were grown to logarithmic phase over 4-5 days at 37°C in Middlebrook 7H9 media with 10% albumin-dextrose-saline, 0.5% glycerol, and 0.05% Tween-80 (“7H9c”). For preparation of aerosol stocks of *M. tuberculosis* H37Rv and Erdman, logarithmic phase cultures were centrifuged at 3,500 rpm for 5 min., resuspended in 1X phosphate-buffered saline (PBS), and sonicated thrice for 30 sec. each at 90% power. Cultures were then centrifuged at 700 rpm for 5 min to pellet large clumps, the supernatant was collected, and then centrifuged again at 700 rpm for 5 min. The supernatant was then collected and the optical density at 600 nM (OD_600_) was measured to determine bacterial concentration, and then aliquoted and frozen at −80°C until use. Frozen stocks of FtLVS (generously gifted by Dr. Karen Elkins, FDA) were cultured in Mueller Hinton Broth containing 1.25 mM CaCl_2_, 2.2 mM MgCl_2_, 0.1% glucose, 0.025% ferric pyrophosphate, and 2% Isovitalex (#211875, Becton Dickinson, Franklin Lakes, NJ) to early log phase (OD600 ≈ 0.25) overnight for use in experiments. For preparation of injection stocks of *F. tularensis* LVS, frozen stocks were thawed at room temperature and then diluted in sterile 1X PBS to the desired concentration.

We used a standardized IC_50_ assay. Briefly, bacteria were grown to an OD_580_ ∼0.5-1, collected by centrifugation, and resuspended in acidified 7H9c. Bacteria were then diluted to an OD580 of 0.02 and 198 µl aliquoted into 30 wells per test condition in 96-well plates. Two-fold dilutions of aldehyde stocks were made and 2 µl of each added in triplicate wells. OD580 was measured in a Molecular Devices M3 plate reader five days later. Data were plotted in GraphPad Prism Plot the data in Prism.

### In Vitro Macrophage Isolation and Culture

Bone-marrow was obtained from *Aldh2^−/−^*, *Aldh2^−/−^Sp140 ^−/−^*, *Sp140 ^−/−^*, *Nos2^−/−^, Nos2^−/−^Aldh2^−/−^, p47phox^−/−^*, p*47phox^−/−^Aldh2^−/−^*, and C57BL/6J mice and cultured in DMEM (#11960-044, Thermo Fischer Scientific, Waltham, MA) containing 10% FBS, 2 mM L-glutamine, and 10% supernatant from 3T3-M-CSF cells (BMDM media) for 6 days with media supplementation on day 3 to generate bone marrow-derived macrophages (BMDMs). BMDMs were then harvested and frozen in liquid nitrogen in 40% FBS, 10% DMSO, and 50% BMDM media until use.

### In Vitro Macrophage Infections

For infection with *M. tuberculosis*, BMDMs were plated in 96-well plates at 5×10^4^ cells/well and allowed to adhere for 48 hours prior to infection. 24 hours prior to infection, media was changed and BMDMs were stimulated with IFNγ (≈ 30 ng/mL) for IFNγ+ conditions. 5 days prior to infection, a frozen stock of *M. tuberculosis* Lux-Erd was thawed and cultured in 7H9 at 37°C to log phase (OD600 = 0.2 – 1.0). *M. tuberculosis* was centrifuged at 3500 rpm for 5 min. and washed twice in PBS, centrifuged at 500 rpm for 5 min. to pellet clumps, and then sonicated at 90% power (twice, 30 sec. each). After sonication, cultures were diluted in DMEM + 5% horse serum + 5% FBS and added to the BMDMs at a multiplicity of infection (MOI) of 1, followed by spinfection at 1200 rpm for 10 min. Supernatant was then removed and the infected BMDMs were washed twice with 1X PBS before the appropriate media was added, and the cells were incubated at 37°C/5% CO_2_ until the first time point. CFU were enumerated at the indicated time points by washing infected cells twice with PBS, lysing in sterile-filtered deionized water with 0.5% Triton X-100 for 10 min. at 37°C/5% CO_2_, and performing serial dilutions in 1X PBS containing 0.05% Tween 80 before aliquoting dilutions onto Middlebrook 7H10 plates supplemented with 0.5% glycerol and 10% OADC (Middlebrook) and incubated for 21 days at 37°C. CFU is normalized to inoculum dose, calculated by plating t = 0 CFU immediately after spinfection. For infection with FtLVS, BMDMs were prepared as described above. BMDMs were plated in 96-well plates at 5×10^4^ cells/well and allowed to adhere for 48 hours prior to infection. 24 hours prior to infection, media was changed and BMDMs were stimulated with IFNγ (≈ 30 ng/mL) for IFNγ positive conditions. To infect the BMDMs, FtLVS stocks were thawed, resuspended in BMDM media, added to the BMDMs at an MOI = 1 and allowed to phagocytose for 2 hours at 37°C/5% CO_2_. After phagocytosis, infected BMDMs were washed thrice with 1X PBS and then supplemented with normal media until predetermined time points. To enumerate CFU/well, BMDMs were rinsed once with 1X PBS, lysed with sterile-filtered deionized water for 10 min., and then serial dilutions were plated on Mueller Hinton plates containing 85 mM NaCl, 1% (w/v) proteose peptone/tryptone, 8% (w/v) bacto-agar, 0.1% glucose, 0.025% ferric pyrophosphate, 2% Isovitalex, and 2.5% bovine serum for 3 days at 37°C/5% CO_2_ until CFU were counted. Cell viability was determined using cell titer glow according to the manufacturer’s protocol. For 4HNE quantification in infected BMDMs, cells were treated as described above and infected via centrifugation at 1200rpm for 10 minutes prior to the generation of cell lysate.

### Aldehyde Quantification

To measure formaldehyde, BMDMs were plated in 96-well plates at 5×10^4^ cells/well and allowed to adhere for 48 hours prior to analysis. 24 hours prior to analysis, BMDMs were stimulated with IFNγ (≈ 30 ng/mL) and PAM3CSK4 (50 ng/mL). BMDM formaldehyde levels were quantified using Mito-RFAP-2, a mitochondrial-targeted version of ratiometric formaldehyde dye RFAP-2 ^50,51^. Briefly, Mito-RFAP-2 was dissolved in DMSO to a concentration of 10 mM and diluted to 10 μM in Hank’s Balanced Salt Solution (HBSS); the probe was then added to the BMDMs for 30 min. at 37°C/5% CO_2_. BMDMs were then rinsed once with HBSS to remove the probe and allowed to incubate in BMDM media for 60 min. at 37°C/5% CO_2_. The cells were then analyzed by fluorescent excitation at 405 nm and 488 nm and emission at 510 nm. Fluorescence was measured using a SpectraMax M3 plate reader (Molecular Devices, San Jose, CA) and reported as the ratiometric value of emission at 405 nm and 488 nm excitations.

BMDM 4HNE levels were quantified by ELISA (abcam, ab287803). Cells were plated at 5×10^5^ cells per/well in 24-well plates and allowed to adhere for 48 hours prior to analysis. 24 hours prior to analysis, BMDMs were stimulated as described above or infected with FtLVS at an MOI of 1. Two hours after infection, cells were washed 1x with warm PBS and then lifted using cold PBS. Cells were then pelleted and lysed, and lysates were processed for 4HNE quantification according to the manufacturer’s protocol. To measure 4HNE in murine tissues, mice were infected with 10^5^ *FtLVS* i.d.. Organs were harvested at 3 dpi and processed for quantification by ELISA according to the manufacturer’s protocol (abcam, ab287803).

### Immunoblotting

BMDMs were prepared as previously described and plated in a 24-well plate at 3×10^5^ cells/well, and infected with *M. tuberculosis* H37Rv according to the previously described protocol. 24 hours post infection, BMDMs were rinsed with 1X PBS and lysed with Laemmli Buffer (#1610737EDU, Bio Rad, Hercules, CA). Samples were heat inactivated/sterilized at 100°C for 20 min. prior to removal from the BSL3 facility. Protein lysates were prepared by heating in 1X SDS for 5 min. at 100°C and then analyzed by SDS-PAGE using 4-20% Criterion TGX Precast Protein Gel (#5678094, Bio Rad), rabbit monoclonal antibody (EPR4494) against ALDH2 (#ab133306, Abcam, Cambridge, MA), and an anti-rabbit HRP-conjugated secondary antibody. Blots were developed with Western Lightning Plus-ECL enhanced chemiluminescence substrate (#NEL103001EA, PerkinElmer, Waltham, MA) and a ChemiDoc MP/475D Imaging System (Bio Rad).

### Bulk and Single Cell RNAseq

Single cell RNA sequencing results were extracted from a previously published dataset^52^ on *M. tuberculosis-*infected B6 mice. For bulk RNAseq, BMDMs were prepared as previously described and plated in a 24-well plate at 3×10^5^ cells/well, and infected with *M. tuberculosis* H37Rv according to the previously described protocol. 24 hours post infection, BMDM media was removed and cells were lysed with TRIzol Reagent (#15596018, Thermo Fisher Scientific). RNA was extraction was performed by adding chloroform and centrifuging samples to make a biphasic mixture containing RNA in an aqueous phase. The RNA-containing phase was then collected and cleaned up using a RNeasy Mini Kit (#74104, Qiagen, Hilden, Germany) with on-column DNase digestion according to the manufacturer’s protocol. The isolated RNA was then quantified and analyzed by a Qubit Quantitation Platform and an Agilent 2100 Bioanalyzer, respectively, at the Functional Genomics Laboratory of the California Institute for Quantitative Biosciences (UC Berkeley) and then submitted to GeneWiz/Azenta Life Sciences (South Plainfield, NJ) for library preparation and sequencing. Each sample was dual indexed using NEBNext multiplex oligomers and then quantity and quality were assessed with a Qubit Broad Range Kit and bioanalyzer, respectively. All samples had a RIN > 9.8 and 100 ng of total RNA was used in library preparation. Samples were then prepared for sequencing using a NEBNext Ultra II RNA-seq with Poly-A selection kit and sequenced on a NovaSeq 6000 S4 (partial lane), with a sequencing depth of 29 million paired-end reads per sample. Illumina sequencing data files were pre-processed with HTStream (v. 1.3.0) to remove PCR duplicates, trim adapters, polyA tails, and N characters, and to remove low quality 3’ and 5’ bases^53^ and were checked for quality with FastQC (v. 0.11.7). Sequences were aligned to the mouse primary assembly GRCm39 from GENCODE^54^ and gene counts were generated using STAR (v. 2.7.10b_alpha_230301^55^).

### Aldehyde susceptibility assays

*M. tuberculosis* H37Rv was cultured as described above. To test susceptibility to various aldehydes, cultures were grown to an optical density at 580 nm (OD_580_) of 0.8-1.0. Bacteria were collected by centrifugation at 3,200 *g* and resuspended in 7H9c acidified to pH 5.5 with 1N HCl and clumped cells were removed with a slow-speed spin at 150 *g*. The supernatant was diluted to an OD_580_ of 0.08. 200 µl of bacteria was added to wells of a 96-well plate; for each condition tested, three wells were prepared. Two μL of 1:100 aldehyde stock solutions were added to each well to obtain the final concentrations indicated in the Figure 2 (A-C). Exposure was carried out for a period of six days. Aldehydes were purchased from the following: 4-hydroxynonenal (4HNE) (# 32100, Cayman Chemical, Ann Arbor, MI), malondialdehyde (MDA) (#63287, Millipore Sigma, Burlington, MA), and formaldehyde (FA) (#15710, Electron Microscopy Sciences, Hatfield, PA) (16% paraformaldehyde solution, stored in aliquots at - 20°C). To generate nitric oxide (NO), sodium nitrite was added to a final concentration of 3 mM. To test the susceptibility of *F. tularensis* LVS to various aldehydes, we followed a previously described protocol^33^. Briefly, *F. tularensis* LVS was plated on Mueller Hinton plates for 3 days at 37°C/5% CO_2_. Multiple colonies colony were then scraped and used to inoculate a liquid culture (to ensure diversity of the culture) that was maintained at 37°C/180 rpm for 12-14 hours. The culture was then washed twice in 1X PBS and resuspended at an OD600 = 0.01 in 1X PBS and aliquoted into a 96-well plate containing various concentrations of 4HNE (#32100, Cayman Chemical Company), or FA (#F1635, Millipore Sigma) and incubated for 2 hours at 37°C/5% CO_2_. After incubation, CFU/well was enumerated for each condition by plating serial dilutions of the culture on Mueller Hinton plates and incubating at 37°C/5% CO_2_ for 3 days.

### In Vivo Pathogen Challenge

Mice were infected with *M. tuberculosis* H37Rv (*Aldh2^−/−^* and B6 mice), or *M. tuberculosis* Erdman (*Aldh2^−/−^* and *Aldh2^−/−^ Sp140^−/−^* mice) *via* the aerosol route. Aerosol infection was performed using a nebulizer and full-body inhalation exposure system (Glas-Col, Terre Haute, IN). Previously prepared aerosol stocks were thawed and 9 mL of culture diluted into sterile-filtered deionized water was loaded into the nebulizer calibrated to deliver ∼25 CFU/mouse, as measured by CFU in the lungs 1 day following infection (data not shown). At predetermined time points, mice were euthanized, and the lungs were mechanically homogenized in 1X PBS containing 0.05% Tween 80 and serial dilutions were plated on 7H10 plates and incubated for 21 days at 37°C until CFU were counted. For mice infected with *F. tularensis* LVS, injection stocks were prepared as previously described and ∼10^5^ CFU were administered intradermally, as measured by plating serial dilutions of the inoculum on Mueller Hinton plates (data not shown). Due to the immunocompromised nature of *Nos2^−/−^* and *Phox-p47^−/−^*mice, experiments involving these genotypes used an inoculum of ∼10^3^ CFU. 3 or 5 dpi, mice were euthanized, and their lungs and spleens were mechanically homogenized in 1X PBS. Serial dilutions were plated on Mueller Hinton plates and incubated for 3 days at 37°C/5% CO_2_ until CFU were counted. For survival studies, mice were infected, monitored daily for weight loss, and euthanized when weight loss exceeded 15% baseline body weight.

### Tissue Processing for Flow Cytometry

Mice were harvested at 3 dpi for flow cytometry analysis and quantification of bacterial burden (CFU). To create single cell suspensions of organs for analysis, lungs were collected into gentleMACS C tubes (Miltenyi Biotec) containing 3 mL of complete RPMI (cRPMI) supplemented with 70 μg/mL Liberase TM (Merck) and 30 μg/mL DNase I (Millipore Sigma). Tissues were mechanically dissociated using the gentleMACS disassociator (lung_01 program) followed by enzymatic digestion for 30 minutes at 37 °C. Samples were then homogenized into single-cell suspensions using the lung_02 gentleMACS program. Digestion was quenched with 2 mL PBS containing 20% fetal bovine serum (Thermo Fisher Scientific), and suspensions were passed through 70 μm cell strainers. Spleens were homogenized by manual dissociation through 70 μm cell strainers.

### Flow Cytometry

For flow cytometry, spleen and lung single-cell suspensions were pelleted and resuspended in 2 mL PBS. A 200 μL aliquot was blocked with BD α-mouse CD16/32 Fc block for 30 minutes at room temperature. Cells were then surface stained for 30 hour at room temperature with optimized dilutions of the following antibodies: BUV805-labeled CD45 (BD 752415), BUV563-labeled Ly6C (BD 755198), BUV496-labeled CD11c (BD 750450), BV786-labeled CD64 (BD 569507), BV605-labeled CD11b (BioLegend 101257), PerCP-Fire806-labeled Siglec-F (BioLegend 155536), FITC-labeled MerTK (BioLegend 151504), PE-Cy7-labeled NK1.1 (BioLegend 108714), APC-Cy7-labeled Ly6G (BioLegend 127623). Cells were then buffer exchanged to PBS then stained with Aqua LIVE/DEAD fixable viability dye (Invitrogen L34957) fo 30 minutes at room temnperature. Stained samples were fixed for 20 minutes at room temperature using Cytofix/Cytoperm (BD Biosciences). For intracellular staining, fixed cells were incubated with PE-labeled iNOS (BioLegend 696806) and APC-labeled Arg-1 (Invitrogen 17-3697-82). To determine absolute cell counts, fluorescent CountBright counting beads (Invitrogen) were added to each sample. Data were acquired on a Symphony A3 flow cytometer (BD Biosciences) and analyzed using FlowJo version 10 (BD Biosciences).

### Cytokine quantification

Mice were infected with 10^5^ FtLVS I.d.. Tissues were harvested at 3 dpi and homogenized using a Cole-Parmer Tissue-Homogenizer (UX-04727-01). Cytokine concentrations in tissue homogenates were quantified using a LEGENDplex™ Mouse Inflammation Panel (Biolegend) per manufacturer’s protocol with volumes adjusted for a 384 well plate format.

### Candidate Gene Analysis Study

Study subjects from the Vietnam human genetics collection and the SCHS have been described in detail previously^56^. In brief, the Vietnamese PTB participants were recruited at district TB units in Ho Chi Minh City, Vietnam as part of larger clinical studies with strictly defined clinical phenotypes^57^. Patients sampled for the genetics study were 18 years or older, HIV negative, had no previous history of TB treatment and were sputum smear positive. The Vietnamese population controls are otherwise healthy adults with primary angle closure glaucoma who have been previously described^58^. DNA from all participants were genome wide genotyped on either the Infinium OmniExpress-24 or OmniExpress-12 array and standard quality control measures and pipelines were applied as described previously^58,59^ (PMID: 40313261). SCHS is a long-term prospective population-based cohort study focused on genetic and environmental determinants of cancer and other diseases in Singapore^60^. From April 1993 to December 1998, the study recruited 63,257 participants of Singaporean Chinese descent, hailing from the Hokkien and Cantonese dialect groups. Starting from April 1994, a random subset of 3% of participants were invited to contribute blood/buccal cell and spot urine samples. In January 2001, the collection of biospecimens was expanded to include all enrolled participants who provided consent and approximately half of all study participants donate blood or buccal samples for genetic studies. PTB cases were identified through linkage with the National Tuberculosis Notification Registry. Diagnosis is primarily based on passive case detection, where individuals seek medical attention for symptoms such as persistent cough, blood-stained sputum, fever, chills, and night sweats. Confirmatory diagnosis involved a positive sputum smear and microbiological culture of *M. tuberculosis*. Most notifications are reported by public hospitals and the Tuberculosis Control Unit and are supplemented by electronic records from the two mycobacterial laboratories in Singapore, ensuring thorough case capture in the national registry. A total of 27,308 samples were whole genome genotyped using the Illumina Global Screening Array (GSA). An additional 2,161 independent subjects from the SCHS CAD-nested case-control study were genotyped on Illumina HumanOmniZhonghua Bead Chip. Comprehensive information regarding genotyping and QC protocols has been previously published^61,62^.

Genotype data for rs671 was extracted from genotyping array data for subsequent statistical analysis. Logistic regression was applied with adjustment for the first three principal components (PCs) to control for any potential population structure. The analysis was conducted separately for the Vietnamese and the SCHS followed by meta-analysis using a fixed-effect inverse-variance weighted (IVW) method. Cochran’s Q (*P_Het_* < 0.05) were evaluated as an indication of heterogeneity. All analysis was conducted using Stata/SE15.1(Statacorp, College Station, TX, USA).

### Statistical Analysis

Data are presented as mean values and error bars represent standard deviation. The number of samples, experimental replicates, and statistical tests are described in the legend of each figure. Statistical analyses were performed using GraphPad Prism 9 (GraphPad, La Jolla, CA) and a *p* < 0.05 was considered significant.

---

**Figure S1.**
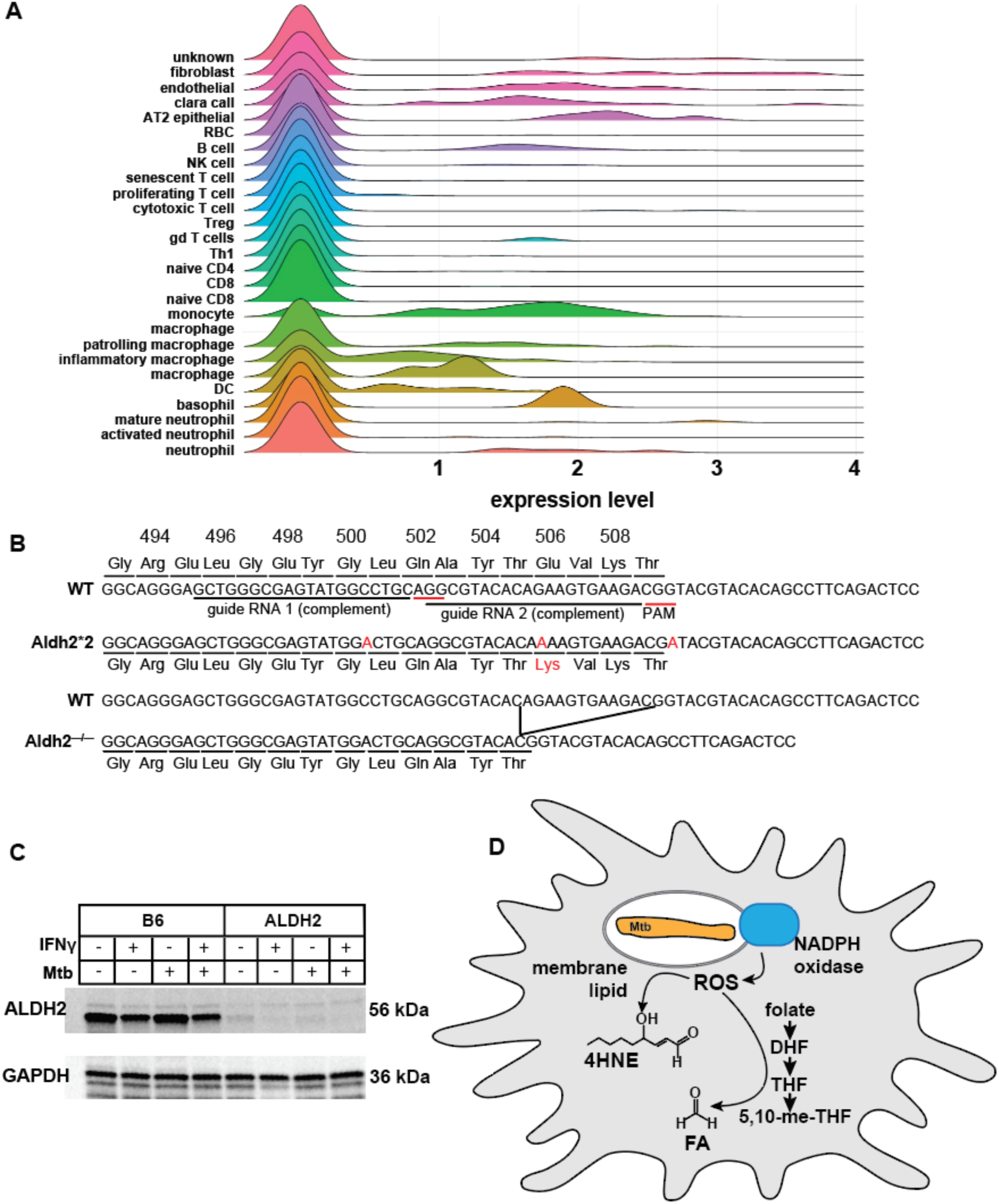
Genetic knock out of *Aldh2* inhibits ALDH2 production. (A) Expression of *Aldh2* in indicated cell types in C57BL/6 mice infected with Mtb for 3 weeks. (B) CRISPR targeting strategy of *Aldh2*2* mice, resulting in mutants containing three adenine point mutations (first and third adenines disrupt sgRNA binding, while second adenine is the human mutation) (*Aldh2*2* mice) and mice containing a 12 bp (four amino acid) in-frame deletion (*Aldh2^−/−^* mice). (C) Immunoblot of WT and *Aldh2^−/−^* BMDM cell lysates stained with αALDH2 (56 kDa) and αGAPDH (36 kDa, control). (D) Model of macrophage sources of 4HNE and FA. 4HNE is derived from peroxidation of host lipids induced in part by activity of reactive oxygen species (ROS) produced by NADPH oxidase acting on cell membranes. Formaldehyde (FA) is derived from decomposition of tetrahydrofolate and its derivatives in a process that is accelerated by ROS.

**Figure S2.**
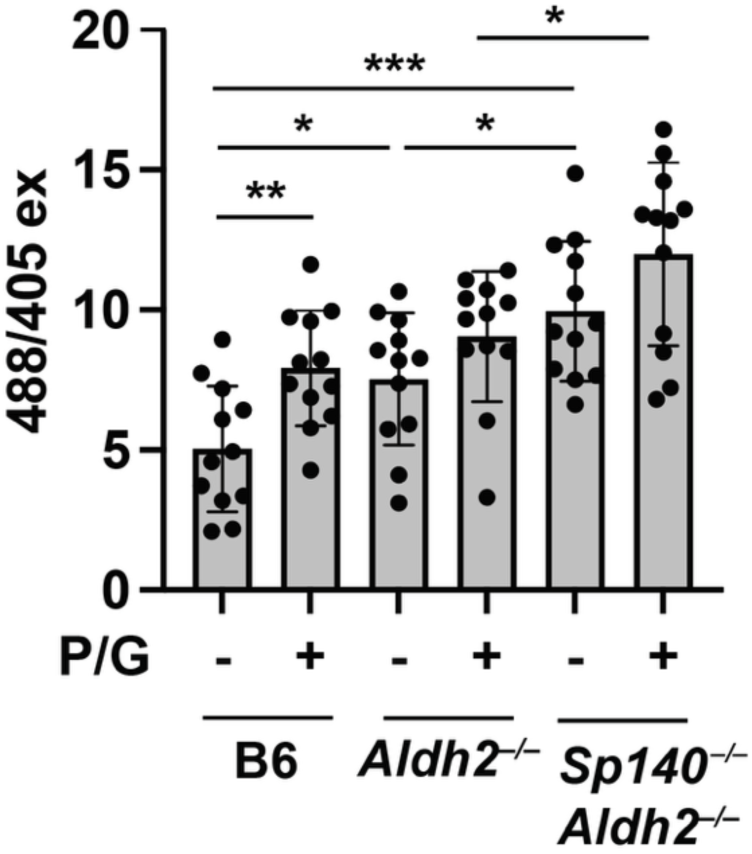
*Sp140^−/−^Aldh2^−/−^* demonstrate increased mitochondrial aldehyde levels. (A) WT, *Aldh2^−/−^*, and *Sp140^−/−^Aldh2^−/−^* BMDM mitochondrial-associated formaldehyde levels after 24 hours stimulation with IFNγ and TLR2-agonist PAM3CSK4 *in vitro* (n = 3 independent experiments, B6 and *Aldh2*^−/−^ results are same as reported in Figure 3A). *, *p* < 0.05, **, *p* < 0.01, ***, *p* < 0.001, Unpaired t-test.

**Figure S3.**
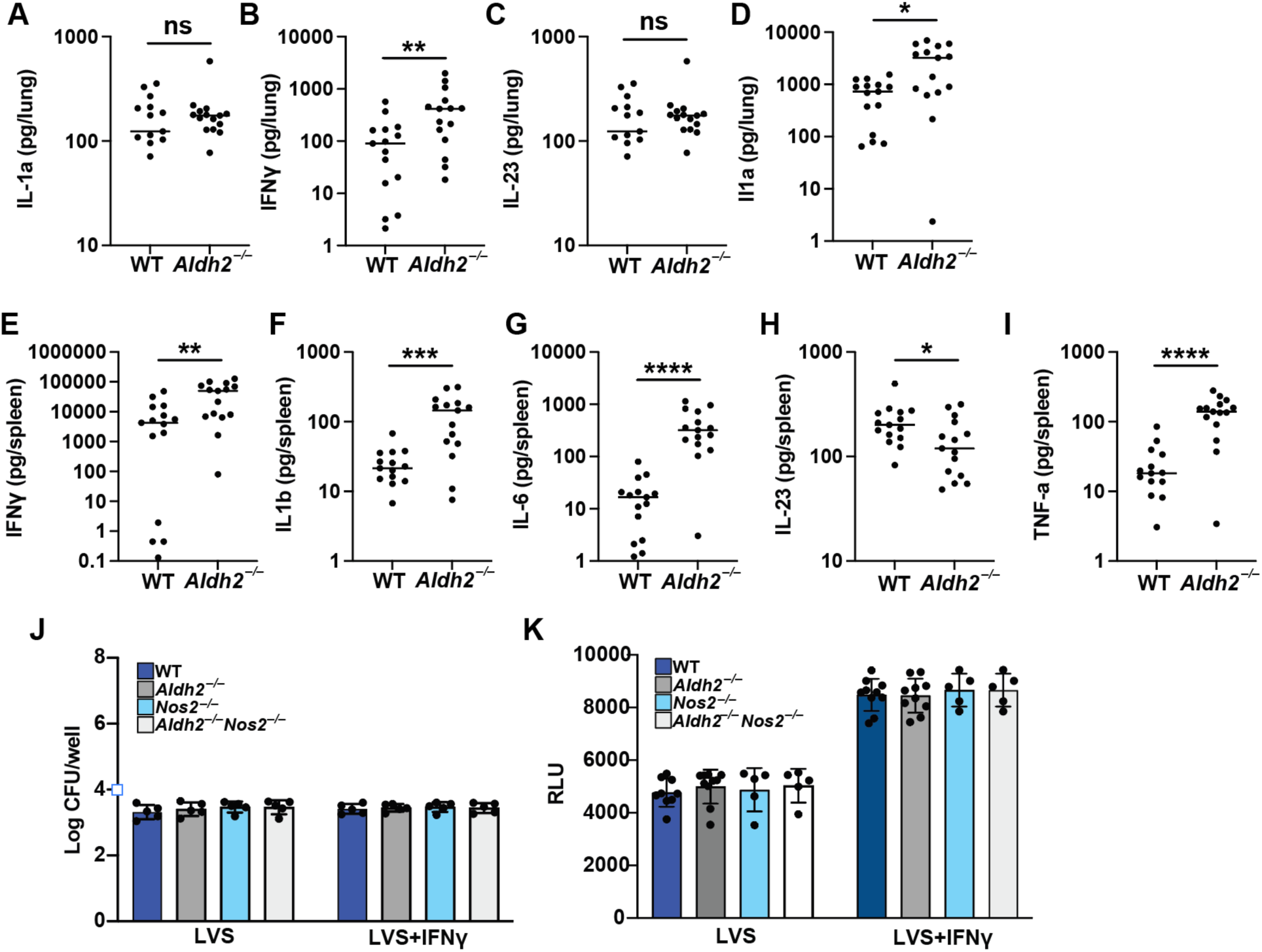
*Aldh2* deficiency confers iNOS-independent resistance to FtLVS with altered lung and spleen cytokines. Quantification of Il-1a (A), IFNγ (B), IL-23 (C), and IL1a (D) in the lungs of WTand *Aldh2^−/−^* mice infected with 10^5^ FtLVS i.d., 3 dpi. Quantification of IFNγ (E), IL1b F), Il-6 (G), IL-23 (H), TNF-a (I) in the spleens of WT and *Aldh2^−/−^*mice infected with 10^5^ FtLVS i.d., 3 dpi. (J) CFU enumeration in WT, *Aldh2^−/−^, Nos2^−/−^,* and *Nos2^−/−^Aldh2^−/−^* infected with FtLVS MOI=1, +/− IFNγ 0 hours post infection. (K) Viability of B6, *Aldh2^−/−^, Nos2^−/−^,* and *Nos2^−/−^Aldh2^−/−^* BMDMs infected with FtLVS MOI=, 1 +/− IFNγ, 2 dpi. (A-I): unpaired t-test. (J-K): 2way ANOVA with Tukey’s multiple comparisons test. Data in (J-K) correspond with results in figure 3G. *p<0.05, **p<0.01, ***p<0.001, ****p<0.0001. Experiments are representative of a minimum of three (J), (K) or three experiments combined (A-I).

**Figure S4.**
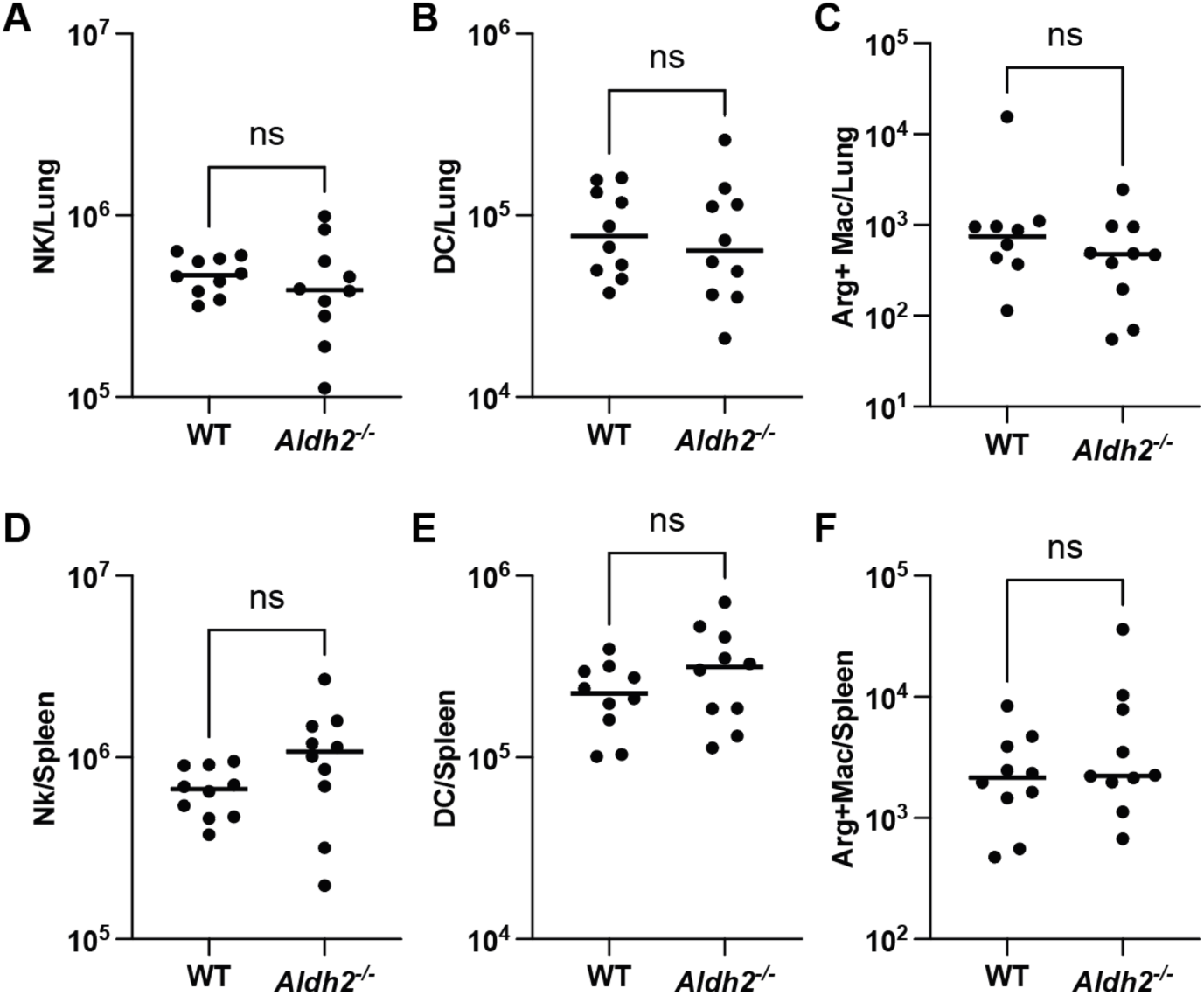
Immune cell frequencies in organs of FtLVS infected, *Aldh2^−/−^* and WT mice. Frequencies of natural killer (NK) cells (A) dendritic cells (B) arginase-positive (Arg+) macrophages (C) in the lungs of WT and *Aldh2^−/−^* mice infected with 10^5^ FtLVS i.d., 3 dpi. Frequencies of natural killer (NK) cells (D) dendritic cells (E) arginase-positive (Arg+) macrophages (F) in the spleens of WT and *Aldh2^−/−^* mice infected with 10^5^ FtLVS i.d., 3 dpi. (A-J) correspond with results in figure 4A-J. *p<0.05, **p<0.01, ***p<0.001, ****p<0.0001 (A-D): unpaired t-test. (E-F): Mann-Whitney test. Experiments are representative of a minimum of three experiments.

